# scRNA-seq reveals novel genetic pathways and sex chromosome regulation in *Tribolium* spermatogenesis

**DOI:** 10.1101/2023.07.18.549532

**Authors:** Michael Robben, Balan Ramesh, Shana Pau, Demetra Meletis, Jacob Luber, Jeffery Demuth

**Author notes:** Authors contributed equally (Michael Robben, Balan Ramesh).

## Abstract

Insights into single cell expression data are generally collected through well conserved biological markers that separate cells into known and unknown populations. Unfortunately for non-model organisms that lack known markers, it is often impossible to partition cells into biologically relevant clusters which hinders analysis into the species. *Tribolium castaneum*, the red flour beetle, lacks known markers for spermatogenesis found in insect species like *Drosophila melanogaster*. Using single cell sequencing data collected from adult beetle testes, we implement a strategy for elucidating biologically meaningful cell populations by using transient expression stage identification markers, weighted principal component leiden clustering. We identify populations that correspond to observable points in sperm differentiation and find species specific markers for each stage. We also develop an innovative method to differentiate diploid from haploid cells based on scRNA-Seq reads and use it to corroborate our predicted demarcation of meiotic cell stages. Our results demonstrate that molecular pathways underlying spermatogenesis in Coleoptera are highly diverged from those in Diptera, relying on several genes with female meiotic pathway annotations. We find that the X chromosome is almost completely silenced throughout pre-meiotic and meiotic cells. Further evidence suggests that machinery homologous to the Drosophila dosage compensation complex (DCC) may mediate escape from meiotic sex chromosome inactivation and postmeiotic reactivation of the X chromosome.

## Introduction

The process of spermatogenesis evolves rapidly at both the anatomical (Fitzpatrick *et al*, 2022) and molecular levels (Murat *et al*, 2023) despite having a core function that is essential for sexual reproduction across metazoans. The rapid pace of evolution is likely driven by several factors including sexual selection and genetic conflict imposed by selfish genetic elements (Kleene, 2005). Additionally, in many male heterogametic species the X and Y chromosomes lack homology across much or all of their length, which imposes challenges for gene regulation and meiotic segregation (Blackmon & Demuth, 2014, 2015). In many species, the inability to pair by homology induces meiotic sex chromosome inactivation (MSCI). MSCI occurs when unsynapsed X and Y chromosomes are silenced during the meiotic stages of spermatogenesis and is thought to be related to a phylogenetically widely conserved surveillance system that protects against aneuploid gametes by condensing unsynapsed chromatin (MSUC) (Turner *et al*, 2005). This has been observed in insects through high-throughput single cell analysis experiments of both *Drosophila melanogaster* and *Anopheles gambiae* (Raz *et al*, 2023; Page *et al*, 2023). This type of analysis provides high level detail of cell specific differentiation in tissue maps but is limited by the need for quality molecular evidence usually associated with well-studied model organisms (Alfieri *et al*, 2022). Challenges arise when characterizing sequenced cells from non-model organisms that share little homology to better studied species, such as the red-flour beetle *Tribolium castaneum*.

While the molecular basis of spermatogenesis is comparatively well studied in classical model organisms like Drosophila (Demarco *et al*, 2014; Siddall & Hime, 2017), this is not well studied in the model beetle, *Tribolium castaneum*. Tribolium serves as a model for the largest eukaryotic order Coleoptera, which harbors many developmental and genomic differences from Drosophila. These differences provide critical points of evolutionary comparison but often also impose challenges for homologous gene and pathway discovery challenging (Pointer *et al*, 2021). Independent fluorescent microscopy studies have implicated the roles of Tra2, Ana1, β2-tubulin, Rad50, and Enolase in beetle sperm development (Shukla & Palli, 2014, 2013; Fishman *et al*, 2017; Khan *et al*, 2021). Although there are only a few studies regarding the molecular basis for spermatogenesis in beetles, the cell biology of *Tribolium* sperm development is relatively well characterized (Dias *et al*, 2015, 2012; Fishman *et al*, 2017). The early sperm cells (spermatogonia) are small, round, and undergo the first mitotic division to give rise to spermatocytes (Fig 1a-d). The post-mitotic spermatocytes are relatively big and are organized in a cyst, but meiosis occurs asynchronously. The secondary spermatocytes, which are found between meiosis I and II, are smaller than primary spermatocytes and have more condensed DNA. Early spermatids formed after meiosis, and a small gap between the acetylated tubulin-labeled axoneme and the nucleus is maintained throughout spermatogenesis. Then, spermatids transition to ellipsis-shaped nuclei with a severely kinked neck region, and as the spermatids mature, the nuclei narrow and twist into an S-shape. Finally, the nuclei straighten in late spermatids, and spermatozoa have needle-shaped nuclei.

**Figure 1.**
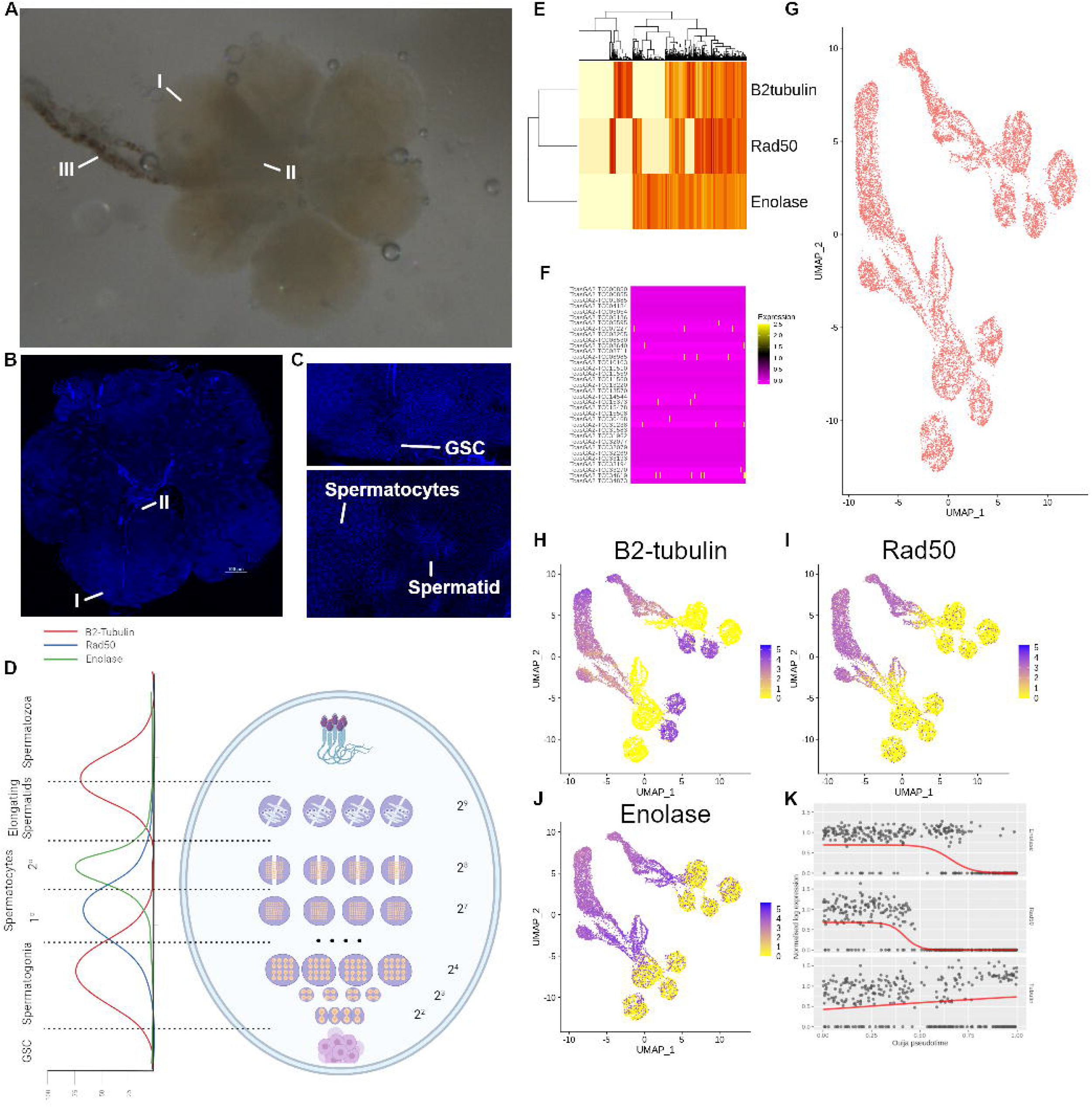
Extraction and single cell sequencing of testes from 4 adult beetles. A,B Microscope pictures of adult beetle testes dissected. (A) Stereomicroscope image of dissected beetle testes in water. (B) DAPI stained slide squash of adult beetle testes. I Distal end of one teste organ II Proximal end of same teste organ III Seminary tubule C Magnified views of cell populations for GSC (upper panel) and Spermatocytes and Spermatids (lower panel) D Diagram of beetle testes with estimated expression levels of previously validated markers; β2-tubulin (TC009035), Rad50 (TC006703), and Enolase (TC011729). Numbers represent the number of cells at each stage of mitotic/meiotic division. Spermatids are represented with 2^9^ cells even though there is no division due to each cyst encompassing 2 bundles of antiparallel sperm. The distal end of testes is at the bottom of diagram and proximal end is at the top. E Expression of previously validated markers for spermatogenic staging across all cells. F Expression of homologues to tissue markers used to stage spermatogenesis in *D. melanogaster* and *A. gambiae* (Table 1). G UMAP projection of weighted principal components showing stage specific clusters. H-J Expression of previously validated stage specific markers in cells displayed as UMAP projection. K Switch-like mechanics of B2-tubulin, Rad50, and Enolase for determining psuedotime status of each cell.

**Table 1.**
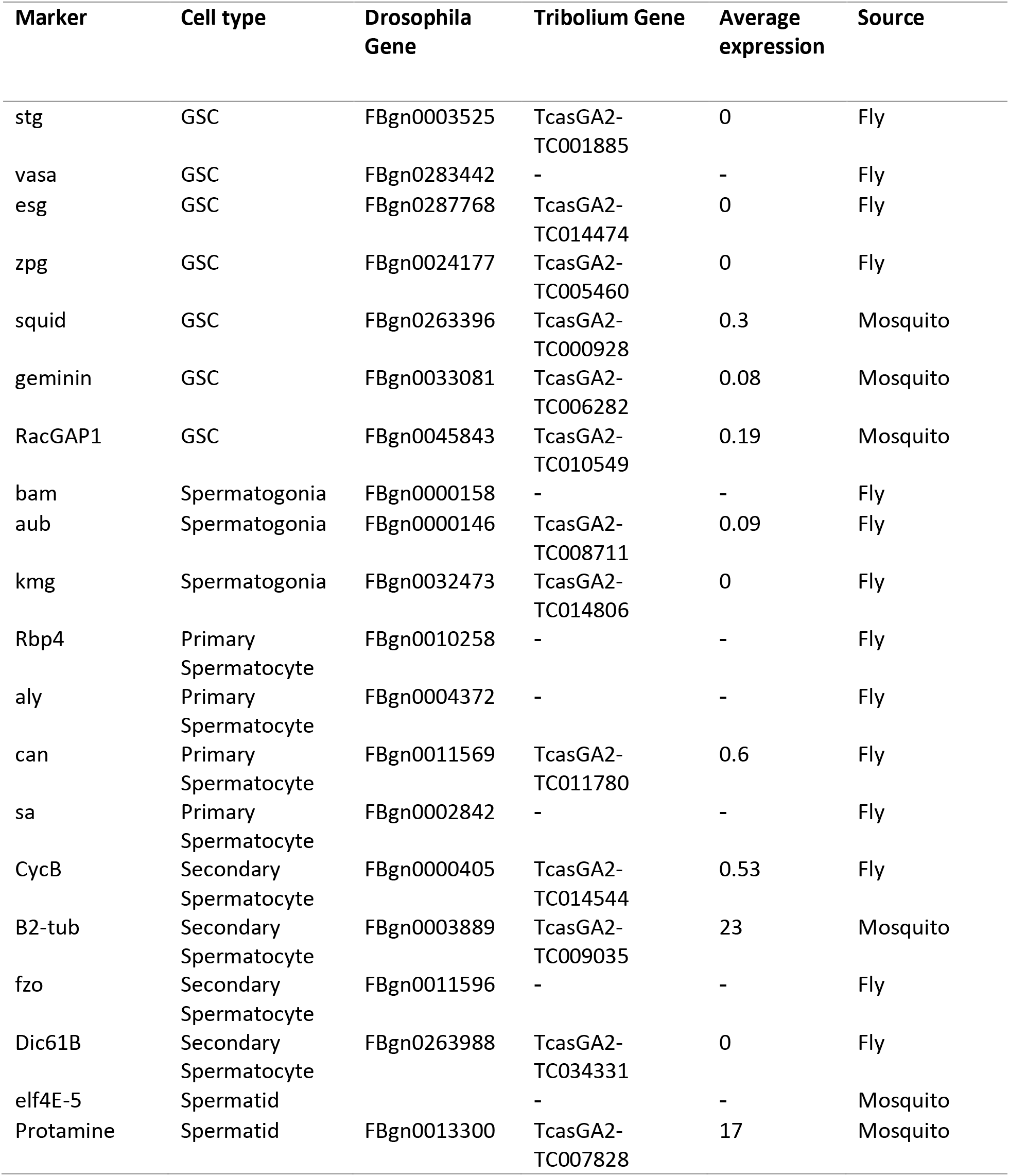
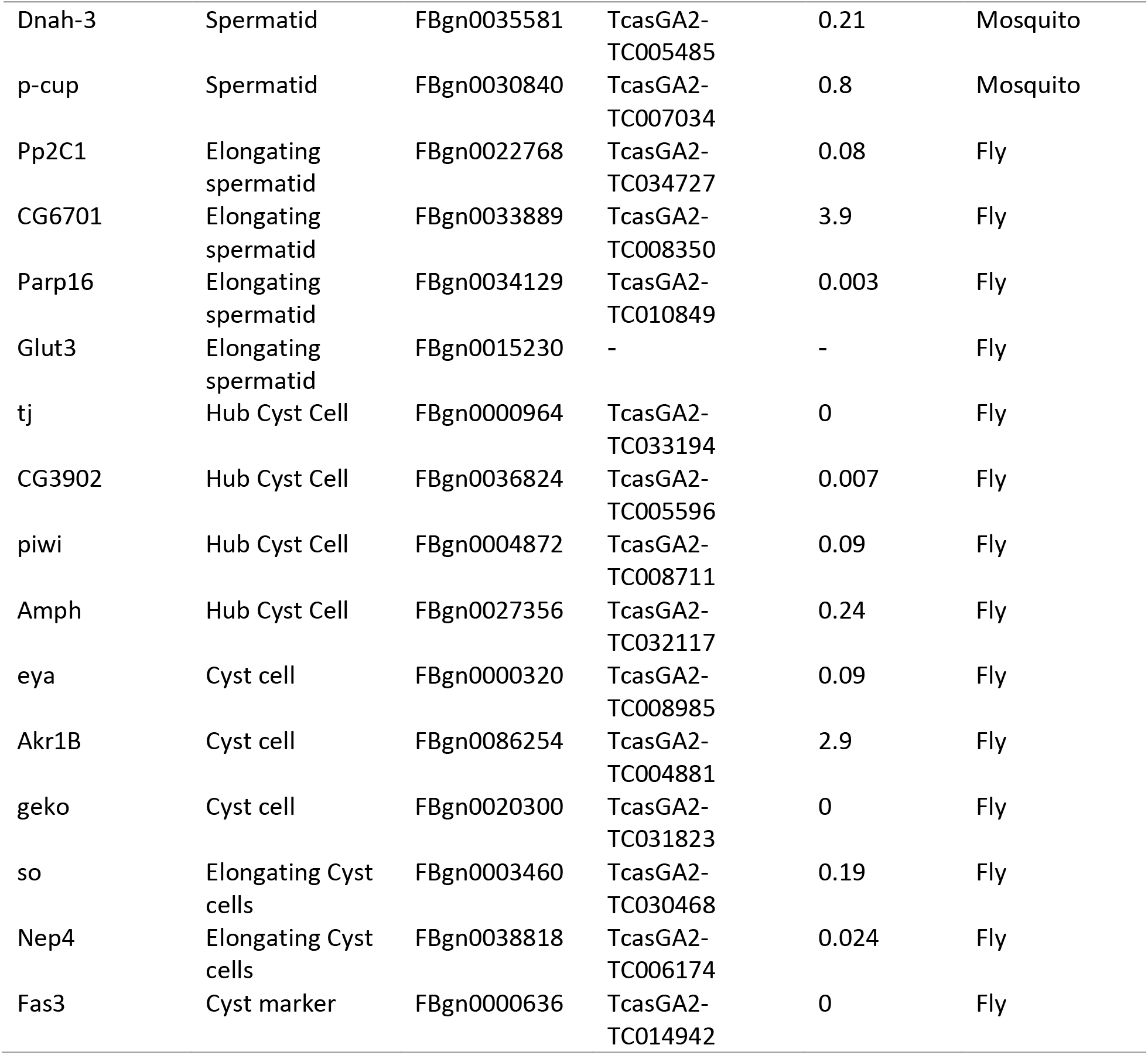
Markers previously used to identify sperm differentiation in insect species. Sources of marker genes from previous scRNA-seq studies in Drosophila and Mosquito (Raz et al, 2023; Page et al, 2023). Average expression refers to normalized mean expression across cells for each marker.

*T. castaneum* uniquely shows interesting dynamics that differentiates it from germ cell development of other insect species. Cyst cells in *Tribolium* encase two antiparallel packets of elongating cells leading to each cyst containing twice the number of flagellated sperms per straw, which has been suggested to develop due to higher competition between males (Fishman et al., 2017). Analyses of sex chromosome gene expression in Tribolium suggest that dosage compensation follows a pattern similar to that of Drosophila, although the molecular mechanisms are less well understood (Prince *et al*, 2010; Mahajan & Bachtrog, 2015; Whittle *et al*, 2020). Furthermore, bulk RNA-seq from male gonads follow a similar pattern to Drosophila wherein X-linked genes appear to be suppressed but not fully silenced (Whittle *et al*, 2020). However, the status of dosage compensation and MSCI in germline tissues remains unresolved because previous studies could not differentiate expression from somatic cells that were included in the gonad samples. To better understand sex chromosome regulation in the Tribolium male germline, we employ high-throughput single-cell transcriptome sequencing from testes-derived heterogeneous cell populations and present an atlas of gene expression throughout spermatogenesis. To compensate for the lack of markers for cell type identification and clustering we develop novel strategies of using non-marker evidence for sperm stage determination.

## Results

### Spermatogenesis Differentiation marker expression

Pooled single cell RNA-sequencing (scRNA-seq) from 4 beetles yielded 1,662,669 cells expressing 16,592 genes. Filtering for lowly expressed genes and cells, doublets, and high mitochondrial expression resulted in 17,069 cells expressing 10,587 features (Supplementary Figure 1a-b). We observed that low variation in gene expression between potential cell groups resulted in poor spatial subdivision within principal component analysis (PCA) and Uniform Manifold Projection (UMAP) (Supplementary Figure c-d). Non-germline or cyst somatic cells were predicted and filtered using common cell line markers (Supplementary Figure 1e-f) leaving 10,587 potential germ line cells and 4,645 possible cyst cells. Due to limited markers for cyst identification, we included cyst cells for further analyses.

Previous reports show that β2-tubulin, Rad-50, and Enolase display well-defined patterns of expression across testis cell types and can be used to stage cells in spermatogenic time (Fig 1e, Supplementary Figure 1g, Kahn et al. 2021). These experimentally validated markers demonstrated more temporal coherence than beetle homologues of Drosophila markers used to previously stage Drosophila testes scRNA-seq data (Fig 1f, Table 1) (Raz *et al*, 2023). Similarly, UMAP projections derived from principal components of the 400 most variably expressed genes did not result in identifiable features of spermatogenesis (i.e., recognizable clusters) or linear relationships to development (Supplementary Figure 1h). This is a common problem in scRNA seq experiments of non-model species that lack identifiable markers of cell type and low variability in gene expression between cells (Alfieri *et al*, 2022).

To resolve cell types in the absence of marker homology with other model systems, we used weighted principal components generated from markers independently shown in other studies to be differentially expressed in germline cell types (Supplemental Table 1) to generate UMAP projections and Leiden clusters. Weighted principal components yielded a higher degree of separation between cell type clusters (Fig 1g, Supplemental Figure 1i). We also observed a greater linear relationship in UMAP space between our 3 marker genes (Fig 1h-j) which we confirmed by psuedotime analysis using Ouija (Fig 1k). Ouija psuedotime displayed higher linearity in UMAP projection from weighted principal components than by those using variable genes alone (Supplementary Figure 1j).

### Tribolium lacks classical marker genes for spermatogenesis

We found little correlation between Drosophila or Anopheles markers of spermatogenesis and marker expression in Tribolium (Table 1, Supplementary Figure 1k-m) (Raz *et al*, 2023; Page *et al*, 2023). This is because essential markers for sperm staging in the two dipterans such as vasa, fzo, or bam (Raz *et al*, 2023), does not seem to play the same role in beetle spermatogenesis. While interesting from an evolutionary perspective, the lack of functional homology provided a challenge for clustering cells by spermatogenic stage. To address this, we chose to look for other, non-expression markers, that could be used to properly stage cell types. Homologues of genes used to phase mitotic cycle in Drosophila were variably expressed across clusters (Fig 2a-b). We found that predicted clusters lined up with markers of mitotic and meiotic activity (Fig 2c) and predicted mitotic phase, with G2M being highest at early mitotic division in Germline Stem Cells (GSC) and spermatogonia and the end of meiosis II in primary spermatocytes.

**Figure 2.**
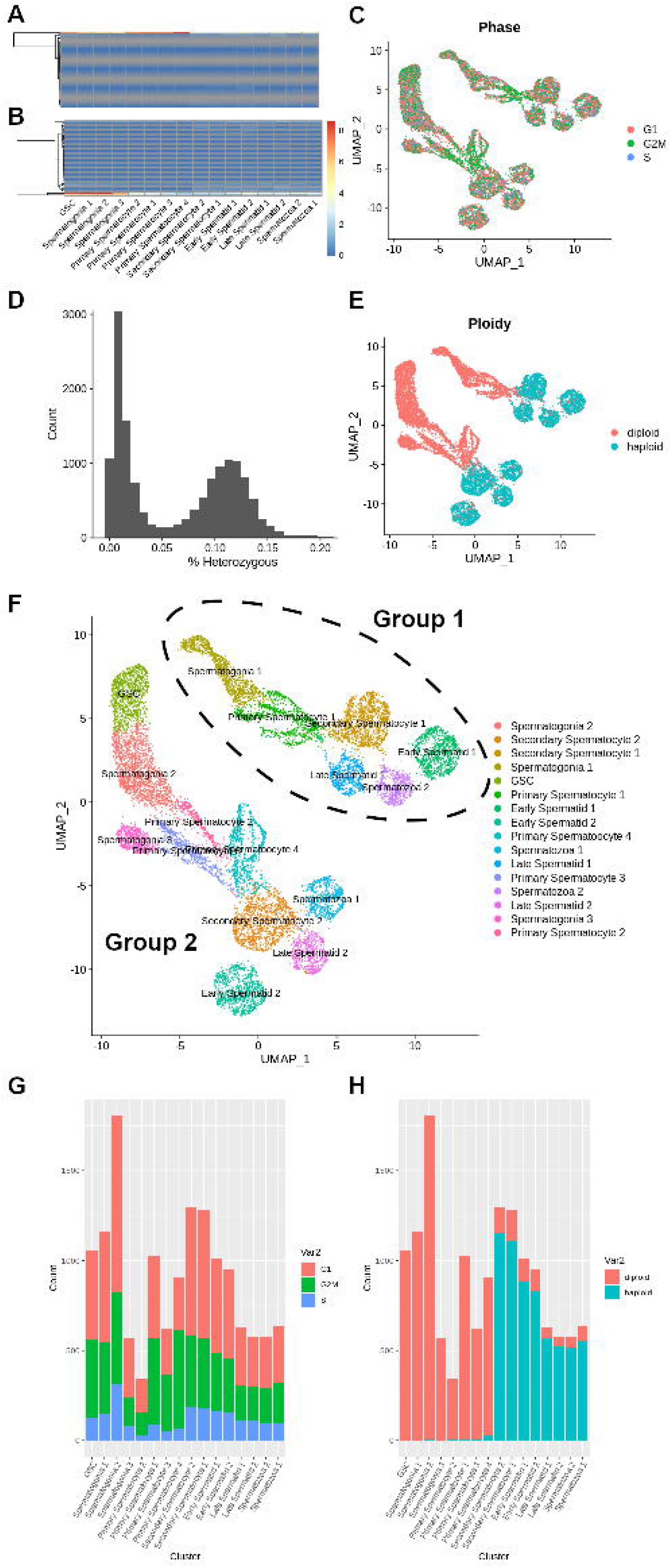
Identification of cluster cell types using phasing evidence. A,B Expression of (A) G2M and (B) S phase specific expression markers. C Estimated mitotic phase of each cell displayed in UMAP projection from homologous Drosophila markers. D Histogram showing the distribution of cells as a function of their percent heterozygosity. Heterozygosity estimated from SNPs in sequenced reads may not represent actual allele status due to low sequencing coverage and variable gene expression. E Ploidy status of each cell as inferred from cells < 5% heterozygous across alleles. F Inferred cell type using multiple evidence. Parallel clusters independently separated during UMAP construction represent 2 distinct groups labeled group 1 and group 2. G,H Stacked barplots showing the total numbers of (G) G1, G2M, and S and (H) haploid and diploid phased cells in each cluster.

We further attempted to computationally validate our cell clustering by differentiating post-meiotic, haploid, from meiotic and pre-meiotic, diploid cell populations. By predicting SNPs from sequenced RNA, we were able to predict the heterozygous content of each cell at all alleles (Fig 2d). Because haploid cells are in theory homozygous, we used a cutoff of 2.5% heterozygosity to phase diploid cells (Fig 2e) which represented secondary spermatocytes, spermatids and spermatozoan cells in our data. In our dataset we found 8,283 diploid cells and 6,180 haploid cells. Using this information, rudimentary labels were given to clusters based on relative values of mitotic, ploidy, and temporal data. Cluster identities were refined and confirmed through differential expression of markers important to stage dependent activity (Supplementary Table 2)

While clusters evenly represented stages of spermatogenesis, we observed two distinct groups of clusters, which we refer to as group 1 and group 2. We found that there were 7,659 cells in group 2 and 6,804 cells in group 1 and there was proportional representation of both clusters and sequenced individuals in both groups (Supplementary Figure 2a, Supplementary Table 3). The relative proportion of cell count in each cell type matched roughly with the cell density of regions of the testes measured by microscopy (Supplementary Figure 2b).

### Non-classical markers of sperm differentiation in Tribolium

While non-expression markers (e.g. ploidy) can confirm rough temporal division in spermatogenesis, we also employed additional methods that provide more context in our analysis. Specifically, psuedotime of our validated marker genes lined up perfectly with UMAP projections and predicted clusters (Fig 3a-b). We also observed linear relationships in the UMAP projections when plotted with psuedotime and RNA velocity values (Fig 3c-d), however the RNA velocity displayed recursive patterns that psuedotime didn’t. We have confirmed, however, that the velocity equation itself does not derive a linear relationship from the specific markers that have been shown to stage spermatogenesis in beetles (Supplementary Figure 2c).

**Figure 3.**
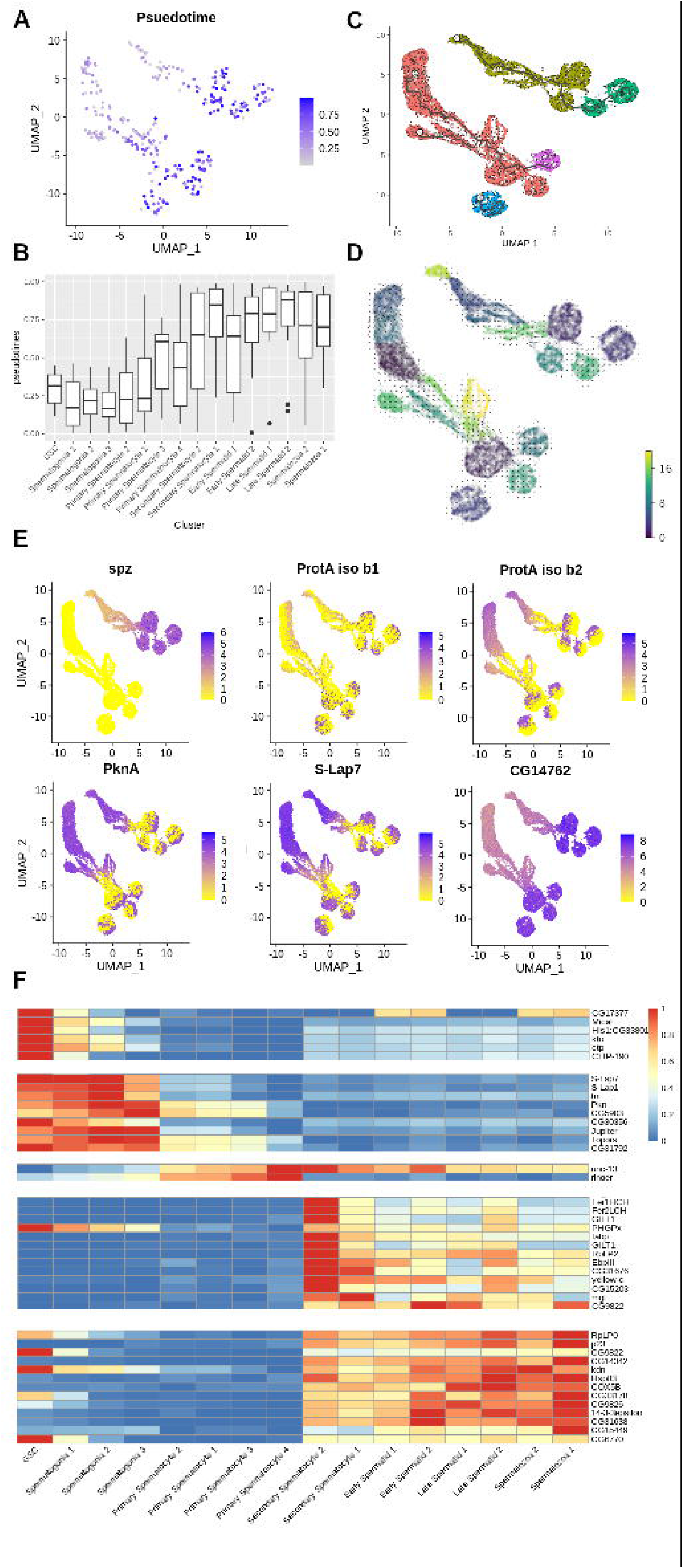
Differentiation trajectory and marker expression of beetle testes. A, B The linear development of sperm cells as inferred by ouija psuedotime analysis of B2-tubulin, Rad50, and Enolase is consistent with labeled clusters. Reported as per cell expression on (A) UMAP projection and (B) per cluster boxplot. C,D Trajectories are also reported as determined by (D) monocle v3 and (D) SCvelo RNA velocity. E Markers that show some differentiation and linear relationships between or within clusters are reported as expressions of each cell in UMAP projection. F Markers identified from monocle derived gene expression markers and represented as normalized average expression values across clusters on a heatmap.

Due to the lack of differentiation markers in beetles that have been identified in other species, it was important to find potential unique markers that could be used to predict cell types in beetle spermatogenesis. Markers that have been shown through molecular evidence to correlate with stages of spermatogenesis, Tra2 and Ana1, were lowly expressed in single cell data (Supplementary Figure 2d-e) (Fishman *et al*, 2017; Shukla & Palli, 2013).

A majority of the highest expressed genes did not show variable patterns of expression across cell clusters, but a few genes were identified that did. The most striking expression pattern we discovered was that of the Tribolium homolog of Spaetzle (spz), which was expressed in group 1 clusters but not group 2 (Fig 3e). Two isoforms of Protamine A (ProtA) exhibited strange expression patterns with isoform b1 showing variable and polar expression within all clusters and groups, while b2 only showed variable expression within secondary spermatocytes, spermatids, and spermatozoa (Fig 3e). These split cluster expression patterns were not opposite between the two isoforms, but we did see opposing expression from isoform b2 in protein kinase A (PknA) (Fig 3e). We also saw similar expression patterns between Sperm-Luecylaminopeptidase 7 (S-Lap7) and ProtA iso b1 (Fig 3e). The Tribolium homolog of CG17377 has an interesting pattern as well and may be a general marker of meiosis as it increases until secondary spermatocytes (Fig 3e). CG17816 is known for negative regulation of microtubule binding and is found to increase expression as it goes into meiosis II (Supplemental Fig 2g) (Rodrigues *et al*, 2022). Unc-13 is interesting as it shows an opposite pattern of expression to that of Drosophila where it builds up to secondary spermatocytes and then decreases in expression in later stages (Supplementary Figure 2g).

Using monocle3, we were able to predict modules of gene co-expression that correlated with stages of spermatogenesis (Supplementary Figure 2h). This allowed us to find markers that could be used to stage cell differentiation in beetle spermatogenesis (Fig 3f). Due to expression of Akb1r, we determined that there was likely equal contamination of cyst cells in germline clusters. While we have little evidence that Akb1r is a marker for cyst cells in beetles we did observe higher median expression of Akb1r in diploid cells than haploid, however the gene would require further validation to confirm that it is a cyst cell marker in beetle (Supplementary Figure 2i-j). We did confirm that the increased weighting of Akb1r in principal components was still not enough to accurately separate cyst and germ clusters and so was not considered in downstream analysis (Supplementary Figure 2k). Similarly, we determined that ProtA, while showing interesting expression dynamics, could not be used to correctly stage cyst cells (Supplementary Figure 2l).

### Preference towards female meiotic pathways throughout spermatogenesis

To better understand the unique dynamics of beetle spermatogenesis, we have performed cell level Gene Set Enrichment Analysis (GSEA) on our dataset. We found several patterns of expression for certain Drosophila Gene Ontology (GO) Biological Processes (BP) terms (Supplementary Figure 3a). There is a stronger enrichment of genes with annotation for female meiotic pathways in contrast to male meiotic pathways (Fig 4a). Spermatogenesis and sperm storage terms increased during spermatogenesis in beetles, but we observed low enrichment of spermatid development genes in late-stage sperm cells (Supplementary Figure 3b). While there was little enrichment of terms for oocyte development, “oocyte dorsal ventral axis specification” was highly enriched, especially in group 1 clusters (Supplementary Figure 3c). Other highly enriched female specific pathways included “imaginal disc derived female genitalia development” and “female gamete generation” (Supplementary Figure 3d). Along with the mitotic phasing of our data, we were also encouraged to see enrichment of “regulation of exit from mitosis” genes be high in GSC and spermatogonia and decrease over time (Supplementary Figure 3e). We also saw similar patterns in GO terms related to germline maintenance (Supplementary Figure 3f).

**Figure 4.**
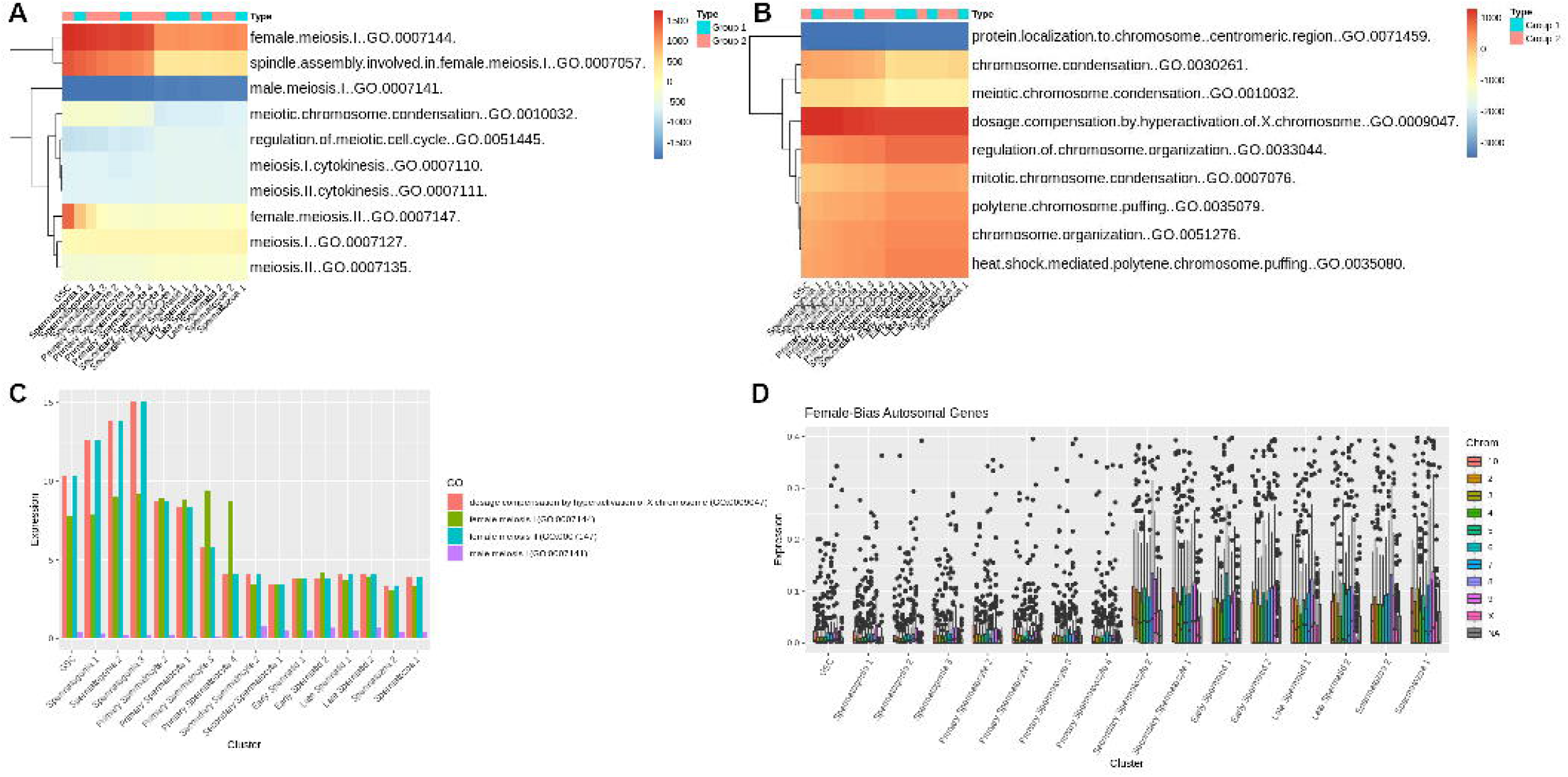
Pathway analysis of cluster expressions. A,B Average enrichment scores per cluster for GO Biological process terms related to (A) Meiosis and (B) Chromosome. Reported values are averaged across per cell enrichment for each term. C Average expression of genes contained within relevant terms. D Expression of female bias genes by cluster by chromosome.

Surprisingly, there was significant enrichment of “dosage compensation by hyperactivation of the X chromosome (GO:0009047)” which we found highly correlated to female meiotic expression (Fig 4b-c). Overall, expression of genes related to female meiosis was greater than those associated with male meiosis (Supplementary Figure 3g-h). When we looked at genes previously associated with male or female biased expression (Prince *et al*, 2010) we found that male biased genes are more strongly expressed in all clusters (Supplementary Figure 4a-b), but the female biased genes had a significant increase in expression in autosomes immediately after meiosis in secondary spermatocytes that we did not see in male biased genes (Fig 4d, Supplementary Figure 4c).

### Incomplete X inactivation during meiotic phase change due to dosage compensation activity

Due to previous studies showing interesting dynamics for X chromosome hyperactivation in somatic tissues and possible MSCI during beetle spermatogenesis (Prince *et al*, 2010), and given the enrichment of hyperactivation pathways that we observed, we further investigated patterns of X-linked gene expression across cell clusters. In pre-meiotic and meiotic cell populations, we observed significantly higher silencing of X-linked genes when compared to autosomal genes than has been found in other species (Supplemental Table 4). However, the mean expression of genes across the X chromosome show selective upregulation at various points during late sperm development (Fig 5a). This appears to correlate with specific regions of the X chromosome that peak in log expression at secondary spermatocyte development (Fig 5b, Supplementary Figure 3d).

**Figure 5.**
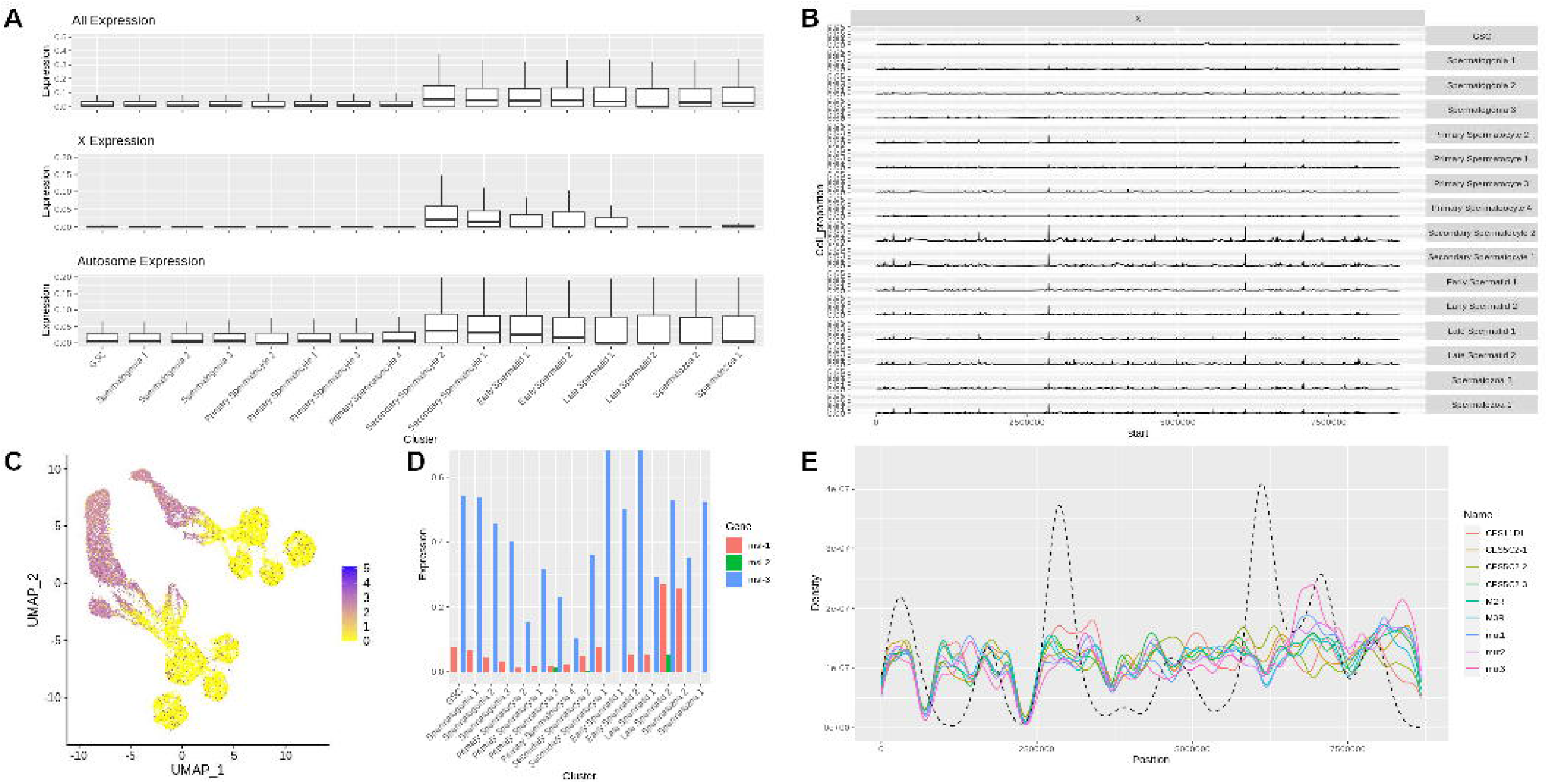
X-inactivation and MSCI of adult beetle testes. A Boxplot showing average expression of each gene across all chromosomes, the X chromosome, and autosomes by cluster. B Percent of cells expressing X chromosome genes above an average of 0.25 normalized expression. C Expression of Unr (TC002472) per cell displayed on UMAP projection D Expression of msl1-3 per cluster. E Density plot showing motif enrichment of DCC CES sites on the X chromosome plotted over mean expression data of X chromosome genes in “Secondary Spermatocyte 1” cluster cells. Dashed line represents density of highly expressed X chromosomal genes

A motif enrichment analysis of upstream promoter regions of the genes that spike in expression through secondary spermatocyte development revealed a motif common to more than half of upregulated genes (Supplementary Figure 4e). Expression of genes associated with the Drosophila dosage compensation complex also increase in expression at similar stages, with expression of male sex lethal 3 (msl-3) and Upstream of N-Ras (Unr) increasing in expression through early stages and msl showing interesting expression dynamics through late spermatid development (Fig 5c-d). It is unknown if roX1/2 expression exists in beetles or if it is also expressed at the same stages. Using previously identified Chromatin Entry Sites (CES) for msl in Drosophila, we predicted DCC occupancy on the X-chromosome (Fig 5e). CES like mut1-3 and CES11D1 showed high linear correlation (p < 0.005) with expression at later stages of sperm development (Supplementary Figure 4f).

## Discussion

A comprehensive analysis of the beetle adult testes through a scRNA-seq atlas provides deep insight into mechanisms of beetle sexual development. In this paper we address a key issue that has prevented similar analyses in non-model organisms like *T. castaneum*, which is the lack of homologous markers used for clustering cells. We show that weighted PCA and non-expression markers can be used to drastically improve variance in dimensionality reduction and properly stage clusters by cell type. Development of these novel methodologies are important as many scRNA-seq analyses rely on the underlying algorithms of clustering packages to form clusters that represent the biological ground truth even though most cannot accurately do this (Krzak *et al*, 2019). We have in our analysis a logical progression of cell differentiation from GSC to Spermatozoa that closely matches the haploid distribution and psuedotime derived from previously validated markers which we know to be accurate to the cell stage. This allowed us to identify new markers that can be used to identify sperm stages in development.

This atlas gives us the ability to directly compare to other studies in Drosophila and mosquito (Raz *et al*, 2023; Page *et al*, 2023). We did not observe separate clusters for germline and cyst cells, likely due to a lack of known beetle markers for cyst cells for the weighted principal components, however the mosquito study also did not identify separate clusters for cyst cells. ProtA and Ak1br might be candidates for germ vs. cyst markers as ProtA replaces histones during sperm development and Ak1br is highly expressed in cyst cells in Drosophila (Okada, 2022; Raz *et al*, 2023). The fact that Tribolium was missing or lacked expression of key Drosophila spermatogenesis genes like fzo, esg, and tj made the overall analysis difficult but highlights the rapid evolutionary dynamics of male gamete development. Many of these genes are shared in mosquitoes which, like fruit flies, belong to the order Diptera (Page *et al*, 2023). The fact that these genes are not expressed in Coleoptera spermatogenesis suggests that critical genes that have been well studied in Drosophila, and suggested to be critical in insect sperm development, may not actually be reflective of the larger class and phylum. Surprisingly, many enriched pathways and expressed genes are associated with female meiotic pathway annotations in other species, indicating that there may be less divergent meiotic pathways between sexes in Coleoptera.

Our results show that X chromosome gene regulation is extremely dynamic and includes MSCI, although a few X-linked genes escape complete inactivation. We observed that expression of X-chromosome genes is already extremely depressed in GSC and rises in expression until secondary spermatocytes. The main difference with other species here is the scale, as we only observed an X-Autosome expression ratio of 0.2% while other insect species note minima never crossing below 20% (Raz *et al*, 2023; Page *et al*, 2023). This implies an overall suppression of the X chromosome and MSCI more absolute than other species and localized entirely to meiotic cells unlike what has been observed in Drosophila (Mikhaylova & Nurminsky, 2011; Meiklejohn *et al*, 2011; Vibranovski *et al*, 2012; Landeen *et al*, 2016; Witt *et al*, 2021; Mahadevaraju *et al*, 2021; Prince *et al*, 2010).

Where it has been studied in detail, MSCI is critical for fertility, however its purpose and the evolutionary factors influencing its origins remain unclear. One hypothesis is that MSCI is driven by sexual antagonism and X inactivation, the so called ‘SAXI hypothesis’ (Wu and Xu, 2003). Since X chromosomes spend 2/3 of their time in females, the X chromosome may accumulate female-beneficial genes even at the expense of those genes’ effects in males. Consequently, MSCI may arise to protect male gametogenesis from these sexually antagonistic effects by silencing X-linked genes during spermatogenesis. Our data are consistent with one aspect of the SAXI hypothesis, that the X chromosome will become demasculinized as male biased and/or testes expressed genes are translocated to autosomes to avoid inactivation (Vibranovski et al., 2009). We found more female-biased genes (186) on the X chromosome than male-biased genes (1). However, SAXI also predicts that there should be a greater concentration of the very early spermatogenic genes on the X chromosome than on autosomes, which we do not see. Overall X expression increases in later stages of Tribolium spermatogenesis including expression of both male and female biased genes.

Alternative hypotheses for the origins of MSCI suggest that it may be a form of host genome defense, protecting against selfish genetic elements, suppressing sex ratio distorters, and /or preventing non-homologous recombination between the X and Y chromosomes (Hamilton, 1967; McKee and Handel, 1993; Meiklejohn and Tao, 2010; Namekawa and Lee, 2009). This line of reasoning follows from MSCI’s ancestral origin being MSUC and fits neatly with our observation of early and extreme X chromosome coupled with the unique meiotic mechanism present in Tribolium. Like many species in the Coleopteran subphylum Polyphaga, Tribolium has a unique form of meiosis wherein the X and Y pair at a distance, held together by a protein scaffold that ensures proper meiotic segregation without synaptic pairing. Under these conditions we might expect MSUC to be active as early as Anaphase I in spermatogenesis (Dutrillaux and Dutrillaux, 2009). To our knowledge, Tribolium is the first species where gene expression in this type of meiosis has been explored.

Perhaps as interesting as insights into the cause of silencing, our analyses may also inform mechanisms of post-meiotic reactivation of X-linked genes. We observe site specificity of increased expression over time, which combined with the correlation of DCC CES enrichment at comparative stages of spermatogenesis, leads us to believe that there is selective upregulation of the X chromosome that may be mediated by the dosage compensation machinery and be necessary for post-meiotic sperm development (Landeen *et al*, 2016; Alekseyenko *et al*, 2008). It is curious that X-expression is increased in the male gamete as beetle male sex determination is connected to the Y chromosome. Reports show that silencing of tcTra2 can result in male XX individuals (Shukla & Palli, 2014). We presume that a lack of dosage compensation could negatively affect the proliferation of male beetles in a population making this observed DCC vital to maintaining the sex ratio. Overall, we can say that beetle MSCI behaves in a different manner than other researched insect species and at least in terms of early sex chromosome silencing is more similar the female heterogametic (ZW) chicken (Namekawa *et al*, 2006; Baarends *et al*, 2005; Turner *et al*, 2005).

## Methods

### Preparation and sequencing of testis single-cell RNA-seq libraries

Testes from 4 male beetles were dissected in cold aerated PBS. The resulting eight testes were placed in 200 ml of lysis buffer (100 ml 0.5% Trypsin EDTA + 100 ml of 4 mg/ml collagenase) in preparation for single cell dissociation as previously described (Mahadevaraju *et al*, 2021). The samples were incubated in the lysis buffer for 15 min in ice with gentle mixing every 30 seconds using a pipette. After incubation, we added 20 µL of 1% FBS (Fetal Bovine Serum) to stop the enzymatic activity by gentle pipetting. The sample was filtered through a 40 mm cell strainer coated with 5µl of 0.04% BSA (Bovine Serum Albumin) solution followed by a 5 min centrifugation at 2000 rpm and 4°C. The resulting cell pellet was resuspended in 15 µl of 0.04% BSA solution before further processing. For cell counting, 5 µl of the single-cell suspension were mixed with 5 µl of the trypan blue dye, and the total cell number and the ratio between live and dead cells were analyzed using an automated cell counter (Bio-Rad Automated Cell Counter TC20™). This method yielded high numbers of single cells (∼5 million live cells) with an average of 70–75% viability. We then submitted cells to the UT Southwestern genomics core for library preparation with the 10X Genomics chromium 3’ kit v3 chemistry. Libraries were then sequenced using 150bp PE on Illumina NovaSeq at the North Texas Genome Center (Arlington, Texas).

### Sequencing data processing

Illumina sequencing data was filtered and aligned to Tcas5.2 (Herndon *et al*, 2020) using cell ranger 7.1.0. Cell ranger output was uploaded to R v4.2.3 (R Core Team, 2017) through Seurat v4.1.0 (Hao *et al*, 2021) into a Seurat object containing 1,662,669 cells with expression data for 16,592 genes. Cells were filtered for low expression, doublets, and high mitochondrial content by setting a cutoff of nFeatures > 100 and < 3,000, and percent mitochondrial features < 5%, resulting in 14,443 cells expressing 10,817 genes. Expression data was normalized using “vst” and likely non-germline or cyst cells were filtered by cells expressing marker genes greater than 0.1 normalized expression (Supplementary Figure 1a-b).

### Marker identification and gene annotation

Tcas5.2 gene protein sequences were aligned to Drosophila r6.52 (dos Santos *et al*, 2015) using blastp 2.5.0 with gap penalty of 5 and word size of 3. Previously documented markers for B2-tubulin, Rad50, and Enolase were found at loci "TcasGA2-TC009035", "TcasGA2-TC006703", and "TcasGA2-TC011729" respectively. Drosophila and mosquito (Data ref: Raz *et al*, 2023; Page *et al*, 2023) markers that have been used to identify cell types of the testes were queried through blast annotation (Table 1). Cyst/germline markers in Drosophila were extracted from previous scRNA-seq data using a Wilcoxon ranked sum test and markers differentially expressed with a p < 0.001 that were represented in more than 10% of cells were used for additional analysis.

### Weighted principal components and clustering

To increase the variance of differentiation markers, we applied weighted principal component analysis (Delchambre, 2015), weighting the selected markers (Supplementary Table 1) 10x against the 400 most variable genes. After generating acceptable UMAP projections, we used Seurat native functions to cluster cells with resolutions from 0.25 to 4. Cell types were determined from a combination of staging markers and differential expression markers identified through the findallmarkers function (Supplementary Table 2).

### Trajectory analysis, co-expression and gene set enrichment

Sequencing data were subjected to different forms of trajectory analysis. B2-tubulin, Rad50, and Enolase were used to generate psuedotime estimates on 300 randomly sampled cells using Ouija (Campbell & Yau, 2019). Alignment bam files were used to generate RNA velocity estimates through SCVelo then plotted in Python v3.7.12 using loom files extracted through R. Seurat data was imported into monocle3 (Trapnell *et al*, 2014) cell data sets and used to generate psuedotime estimates of cell differentiation. Using monocle3 functions, we calculated modules of similarly expressing genes using the K-Nearest Neighbor graph method. To determine patterns of expression related to function we turned to gene set enrichment analysis (GSEA) using a gene set constructed from Drosophila Biological Process GO annotations (https://maayanlab.cloud/FlyEnrichr/#stats, downloaded 6/21/2023). GSEA on individual cells was conducted using the escape package (Borcherding *et al*, 2021)in R with 1000 groups and a minimum size of 5. Median enrichment scores were determined per cluster and displayed for most variably enriched terms as well as terms that contained “meiosis”, “mitosis”, “male”, “female”, “sperm”, “cyst”, “germ”, and “chromosome”.

### Ploidy analysis

To determine the fractions of cells that were haploid and diploid in our data set, we sought to estimate the level of heterozygosity in samples through Single Nucleotide Polymorphism (SNP) analysis of the single reads. This can be represented with the equation:

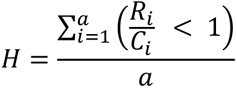

Where H is the percent heterozygosity for each cell, *a* is the total number of alleles for which expression was detected in each cell, *R* is the number of reads mapped to the reference allele, and *C* is the total number of reads at each allele. We predicted SNPs using bcftools mpileup (Li, 2011) to get biallelic markers. Using the program scAlleleCount (https://github.com/barkasn/scAlleleCount) we calculated for each cell the number of reads that mapped to either allele. We then calculated the frequency of reads at each allele site to determine the heterozygosity of each allele in every cell. This allowed us to determine the percent heterozygosity across all detected alleles for each cell.

### Motif enrichment

Previously identified msl CES motifs (Alekseyenko *et al*, 2008) were predicted on the X-chromosome using the Fimo tool in the meme-suite (https://meme-suite.org/meme/index.html). Novel motifs were identified from 1,000 bp regions upstream of highly expressed X-linked genes. Distance was calculated from the start of each motif to the start of each gene and the shortest distance was tested for linear correlation with expression of each gene using a 3-way Anova with TukeyHSD.

## Acknowledgements

We would like to thank Heath Blackmon for graciously providing us the protocol for testes dissection using the Tribolium beetles. We thank the Oliver lab (NIDDK) for providing us prepublication protocols for preparing single-cell suspension from Drosophila testes and the Betran lab at UTA for graciously guiding us with the use the fluorescence microscopy. We are grateful to Beena Margabandhu and Chris Lander, who helped finetune the protocol for the testes dissections in *T. castaneum*. We would also like to thank Avishek Das for contributing code and data processing work for this analysis. This work was funded by the UTA Phi Sigma Biology Graduate Student Society (BR), a UTA Research Enhancement Program award (JPD), the Cancer Prevention Research Institute of Texas (JML), and a Rising STAR’s awards (JML). We thank the Texas Advanced Computing Cluster for the use of their high-performance computing resources.

## Author Contributions

The experiment was designed by MR, BR, JL, and JD. Material was collected, imaged, and sequenced by BR and SP. Computational and statistical analysis was performed by MR and DM. Manuscript was prepared by MR, JL, and JD.

## Data Availability

All code used to generate images and analysis in the manuscript can be found hosted on github: https://github.com/RobbenUTA/BeetleTestes. Sequencing and count data have been uploaded for public access at NCBI SRA and GEO databases under the bioproject accession number PRJNA994887.

## Supplemental Figures and Tables

**Supplemental Table 1.** The markers that were used to construct weighted principal components.

**Supplemental Table 2.** Output of findallmarkers for each cluster.

**Supplemental Table 3.** Number of cells per individual per cluster as predicted by allelic specificity.

**Supplemental Table 4.** Comparison of the mean expression of X-chromosomal and autosomal genes. Significance determined by signed Wilcoxon ranked sum test and adjusted using a benjamini-hochberg false discovery rate.

**Supplemental Figure 1.**

A,B Sequencing statistics for 4 pooled individuals. (A) Total sequenced RNA count by percent mitochondrial expression and (B) total feature count by number of expressed features.

C,D Projections derived from 400 most variable genes with (C) PCA and (D) UMAP.

E Total predicted numbers of various somatic cell types as determined by greater than 0.1 average normalized counts of potential marker genes.

F Total predicted numbers of somatic, cyst, and germ cells. Cyst cells are included in final clustering’s due to lack of validated markers for separation.

G Venn diagram showing the number of cells expressing 3 validated marker genes above a normalized count of 0.1.

H Expression of 3 validated marker genes in UMAP space generated from 200 most variable gene PCA.

I Ouija psuedotime from 3 validated marker genes mapped in UMAP space generated from 200 most variable gene PCA.

J Principal components 1 and 2 from weighted PCA of selected marker genes and 400 most variable genes.

K,L,M Expression of (L) germline and (M) cyst marker genes.

**Supplemental Figure 2.**

A Cell density of testes predicted from DAPI stained microscope images. Cell count estimated from watershed segmentations determined in a 240×72 pixel region at multiple locations from distal to proximal along teste axis.

B Numbers of cells per cluster in scRNA-seq data.

C Linear approximations of RNA velocity across all cells based on expression of 3 previously validated markers.

D,E Expression of (D) Tra2 (TC034297) and (E) Ana1 (TC033159) on UMAP projection.

F,G Expression of (F) CG17618 and (G) Unc-13 homologue on UMAP projection

H Average expression of monocle3 module genes by cluster.

I Expression of Akr1b on UMAP projection.

J Expression of Akr1b in haploid and diploid cells.

K UMAP derived from weighted PCA with increased weight (20x) applied to Akr1b. Each cell is colored by original clustering.

L Expression of ProtA isoforms b1 (TC007670; left panels) and b2 (TC007828; right panels) by cluster in UMAP projections (top panels) and as a boxplot (bottom panels)

**Supplemental Figure 3.**

GSEA enrichment scores plotted on heatmap by cluster. Gene sets derived from drosophila BP of homologous beetle genes. Group of each cluster represented in column annotation.

A The top 100 most variably enriched GO terms by cluster.

B Enrichment of GO terms associated with “sperm” by cluster.

C Enrichment of GO terms associated with “oocyte” by cluster.

D Enrichment of GO terms associated with “male” by cluster.

E Enrichment of GO terms associated with “mitosis” by cluster.

F Enrichment of GO terms associated with “germ” by cluster.

G Expression of genes in “female meiosis” BP gene sets on UMAP projection.

H Expression of genes in “male meiosis” BP gene sets on UMAP projection.

**Supplemental Figure 4.**

A,B Overall expression of (A) male and (B) female-biased genes by cluster shown by heatmap.

B Boxplot showing median expression of male biased genes by chromosome and cluster.

C Mean expression of X-linked genes across clusters.

D Enrichment of motif associated with region 1,000 bp upstream of all highly expressed x-linked genes.

E Correlation of distance to expression between each putative CES by cluster.

## References

Alekseyenko AA, Peng S, Larschan E, Gorchakov AA, Lee O-K, Kharchenko P, McGrath SD, Wang CI, Mardis ER, Park PJ, et al (2008) A sequence motif within chromatin entry sites directs MSL establishment on the Drosophila X chromosome. Cell 134: 599–609

Alfieri JM, Wang G, Jonika MM, Gill CA, Blackmon H & Athrey GN (2022) A Primer for Single-Cell Sequencing in Non-Model Organisms. Genes 13

Baarends WM, Wassenaar E, van der Laan R, Hoogerbrugge J, Sleddens-Linkels E, Hoeijmakers JHJ, de Boer P & Grootegoed JA (2005) Silencing of unpaired chromatin and histone H2A ubiquitination in mammalian meiosis. Mol Cell Biol 25: 1041–1053

Blackmon H & Demuth JP (2014) Estimating tempo and mode of Y chromosome turnover: explaining Y chromosome loss with the fragile Y hypothesis. Genetics 197: 561–572

Blackmon H & Demuth JP (2015) The fragile Y hypothesis: Y chromosome aneuploidy as a selective pressure in sex chromosome and meiotic mechanism evolution. Bioessays 37: 942–950

Borcherding N, Vishwakarma A, Voigt AP, Bellizzi A, Kaplan J, Nepple K, Salem AK, Jenkins RW, Zakharia Y & Zhang W (2021) Mapping the immune environment in clear cell renal carcinoma by single-cell genomics. Commun Biol 4: 122

Campbell KR & Yau C (2019) A descriptive marker gene approach to single-cell pseudotime inference. Bioinformatics 35: 28–35

Delchambre L (2015) Weighted principal component analysis: a weighted covariance eigendecomposition approach. Mon Not R Astron Soc 446: 3545–3555

Demarco RS, Eikenes ÅH, Haglund K & Jones DL (2014) Investigating spermatogenesis in Drosophila melanogaster. Methods 68: 218–227

Dias G, Lino-Neto J, Mercati D & Dallai R (2015) The sperm ultrastructure and spermiogenesis of Tribolium castaneum (Coleoptera: Tenebrionidae) with evidence of cyst degeneration. Micron 73: 21–27

Dias G, Yotoko KSC, Gomes LF & Lino-Neto J (2012) Uncommon formation of two antiparallel sperm bundles per cyst in tenebrionid beetles (Coleoptera). Naturwissenschaften 99: 773–777

Fishman EL, Jo K, Ha A, Royfman R, Zinn A, Krishnamurthy M & Avidor-Reiss T (2017) Atypical centrioles are present in Tribolium sperm. Open Biol 7

Fitzpatrick JL, Kahrl AF & Snook RR (2022) SpermTree, a species-level database of sperm morphology spanning the animal tree of life. Sci Data 9: 30

Hao Y, Hao S, Andersen-Nissen E, Mauck WM 3rd, Zheng S, Butler A, Lee MJ, Wilk AJ, Darby C, Zager M, et al (2021) Integrated analysis of multimodal single-cell data. Cell 184: 3573–3587.e29

Herndon N, Shelton J, Gerischer L, Ioannidis P, Ninova M, Dönitz J, Waterhouse RM, Liang C, Damm C, Siemanowski J, et al (2020) Enhanced genome assembly and a new official gene set for Tribolium castaneum. BMC Genomics 21: 47

Khan SA, Jakes E, Myles KM & Adelman ZN (2021) The β2Tubulin, Rad50-ATPase and enolase cis-regulatory regions mediate male germline expression in Tribolium castaneum. Sci Rep 11: 18131

Kleene KC (2005) Sexual selection, genetic conflict, selfish genes, and the atypical patterns of gene expression in spermatogenic cells. Dev Biol 277: 16–26

Krzak M, Raykov Y, Boukouvalas A, Cutillo L & Angelini C (2019) Benchmark and Parameter Sensitivity Analysis of Single-Cell RNA Sequencing Clustering Methods. Front Genet 10: 1253

Landeen EL, Muirhead CA, Wright L, Meiklejohn CD & Presgraves DC (2016) Sex Chromosome-wide Transcriptional Suppression and Compensatory Cis-Regulatory Evolution Mediate Gene Expression in the Drosophila Male Germline. PLoS Biol 14: e1002499

Li H (2011) A statistical framework for SNP calling, mutation discovery, association mapping and population genetical parameter estimation from sequencing data. Bioinformatics 27: 2987–2993

Mahadevaraju S, Fear JM, Akeju M, Galletta BJ, Pinheiro MMLS, Avelino CC, Cabral-de-Mello DC, Conlon K, Dell’Orso S, Demere Z, et al (2021) Dynamic sex chromosome expression in Drosophila male germ cells. Nat Commun 12: 892

Mahajan S & Bachtrog D (2015) Partial dosage compensation in Strepsiptera, a sister group of beetles. Genome Biol Evol 7: 591–600

McKee BD & Handel MA (1993) Sex chromosomes, recombination, and chromatin conformation. Chromosoma 102: 71–80

Meiklejohn CD, Landeen EL, Cook JM, Kingan SB & Presgraves DC (2011) Sex chromosome-specific regulation in the Drosophila male germline but little evidence for chromosomal dosage compensation or meiotic inactivation. PLoS Biol 9: e1001126

Mikhaylova LM & Nurminsky DI (2011) Lack of global meiotic sex chromosome inactivation, and paucity of tissue-specific gene expression on the Drosophila X chromosome. BMC Biol 9: 29

Murat F, Mbengue N, Winge SB, Trefzer T, Leushkin E, Sepp M, Cardoso-Moreira M, Schmidt J, Schneider C, Mößinger K, et al (2023) The molecular evolution of spermatogenesis across mammals. Nature 613: 308–316

Namekawa SH, Park PJ, Zhang L-F, Shima JE, McCarrey JR, Griswold MD & Lee JT (2006) Postmeiotic sex chromatin in the male germline of mice. Curr Biol 16: 660–667

Okada Y (2022) Sperm chromatin condensation: epigenetic mechanisms to compact the genome and spatiotemporal regulation from inside and outside the nucleus. Genes Genet Syst 97: 41–53

Page N, Taxiarchi C, Tonge D, Chesters E, Kuburic J, Game L, Nolan T & Galizi R (2023) Single-cell profiling of mosquito spermatogenesis defines the onset of meiotic silencing and pre-meiotic overexpression of the X chromosome.

Pointer MD, Gage MJG & Spurgin LG (2021) Tribolium beetles as a model system in evolution and ecology. Heredity 126: 869–883

Prince EG, Kirkland D & Demuth JP (2010) Hyperexpression of the X chromosome in both sexes results in extensive female bias of X-linked genes in the flour beetle. Genome Biol Evol 2: 336–346

Raz AA, Vida GS, Stern SR, Mahadevaraju S, Fingerhut JM, Viveiros JM, Pal S, Grey JR, Grace MR, Berry CW, et al (2023) Emergent dynamics of adult stem cell lineages from single nucleus and single cell RNA-Seq of Drosophila testes. Elife 12

Raz AA, Vida GS, Stern SR, Mahadevaraju S, Fingerhut JM, Viveiros JM, Pal S, Grey JR, Grace MR, Berry CW, et al (2023) ASAP single cell dataset visualization (https://asap.epfl.ch/projects/ASAP24). [DATASET]

R Core Team (2017) R: A Language and Environment for Statistical Computing. (https://www.R-project.org/) [PREPRINT]

Rodrigues LR, Zwoinska MK, Wiberg RAW & Snook RR (2022) The genetic basis and adult reproductive consequences of developmental thermal plasticity. J Anim Ecol 91: 1119–1134

dos Santos G, Schroeder AJ, Goodman JL, Strelets VB, Crosby MA, Thurmond J, Emmert DB, Gelbart WM & FlyBase Consortium (2015) FlyBase: introduction of the Drosophila melanogaster Release 6 reference genome assembly and large-scale migration of genome annotations. Nucleic Acids Res 43: D690–7

Shukla JN & Palli SR (2013) Tribolium castaneum Transformer-2 regulates sex determination and development in both males and females. Insect Biochem Mol Biol 43: 1125–1132

Shukla JN & Palli SR (2014) Production of all female progeny: evidence for the presence of the male sex determination factor on the Y chromosome. J Exp Biol 217: 1653–1655

Siddall NA & Hime GR (2017) A Drosophila toolkit for defining gene function in spermatogenesis. Reproduction 153: R121–R132

Trapnell C, Cacchiarelli D, Grimsby J, Pokharel P, Li S, Morse M, Lennon NJ, Livak KJ, Mikkelsen TS & Rinn JL (2014) The dynamics and regulators of cell fate decisions are revealed by pseudotemporal ordering of single cells. Nat Biotechnol 32: 381–386

Turner JMA, Mahadevaiah SK, Fernandez-Capetillo O, Nussenzweig A, Xu X, Deng C-X & Burgoyne PS (2005) Silencing of unsynapsed meiotic chromosomes in the mouse. Nat Genet 37: 41–47

Vibranovski MD, Zhang YE, Kemkemer C, Lopes HF, Karr TL & Long M (2012) Re-analysis of the larval testis data on meiotic sex chromosome inactivation revealed evidence for tissue-specific gene expression related to the Drosophila X chromosome. BMC Biol 10: 49; author reply 50

Whittle CA, Kulkarni A & Extavour CG (2020) Absence of a Faster-X Effect in Beetles (Tribolium, Coleoptera). G3 10: 1125–1136

Witt E, Shao Z, Hu C, Krause HM & Zhao L (2021) Single-cell RNA-sequencing reveals pre-meiotic X-chromosome dosage compensation in Drosophila testis. PLoS Genet 17: e1009728

Wu CI & Xu EY (2003) Sexual antagonism and X inactivation--the SAXI hypothesis. Trends Genet 19: 243–247

