## Supplementary figures and images for "scRNA-seq reveals novel genetic pathways and sex chromosome regulation in *Tribolium* spermatogenesis"

### SupplementaryFigure1.png

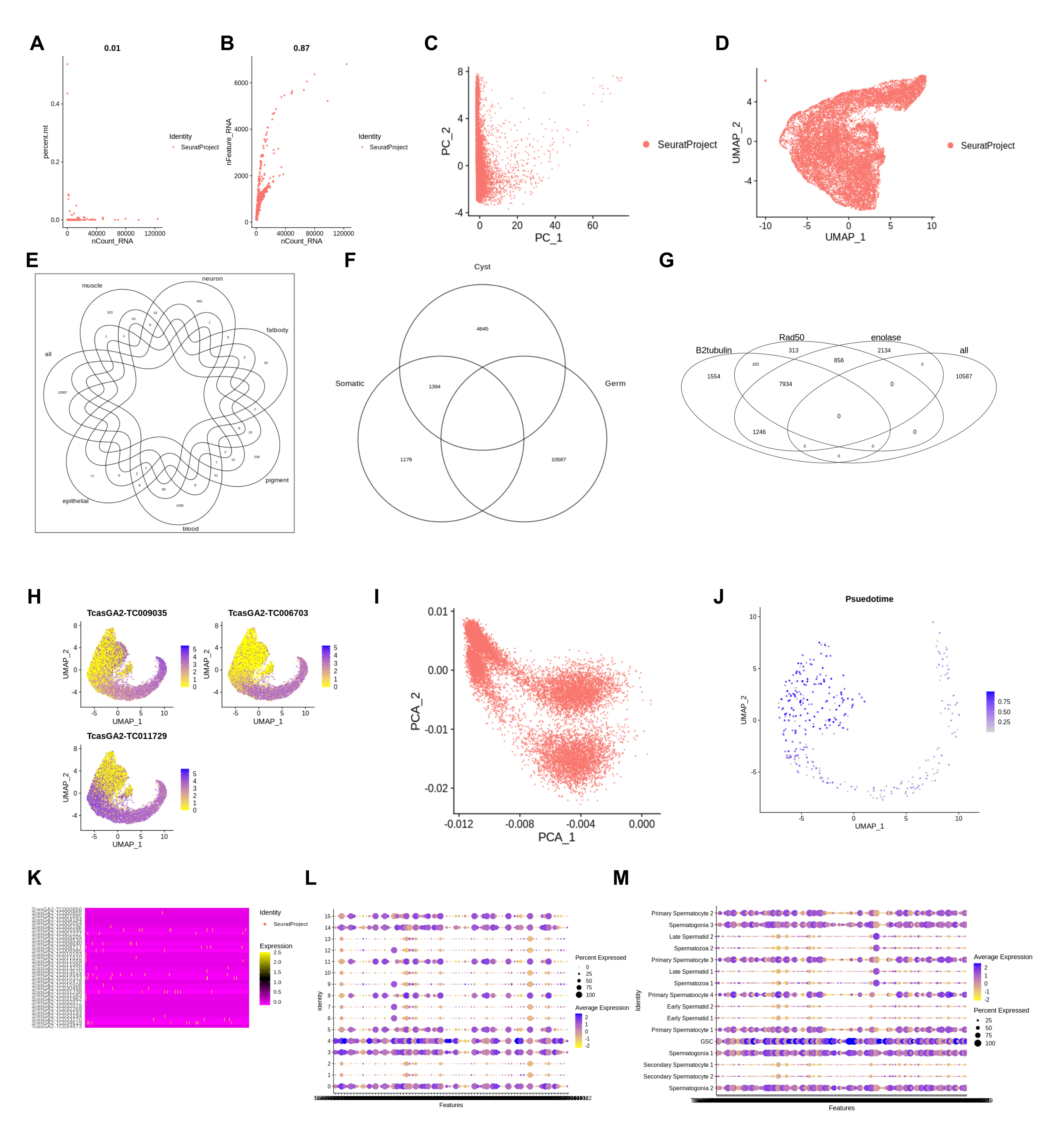

### SupplementaryFigure2.png

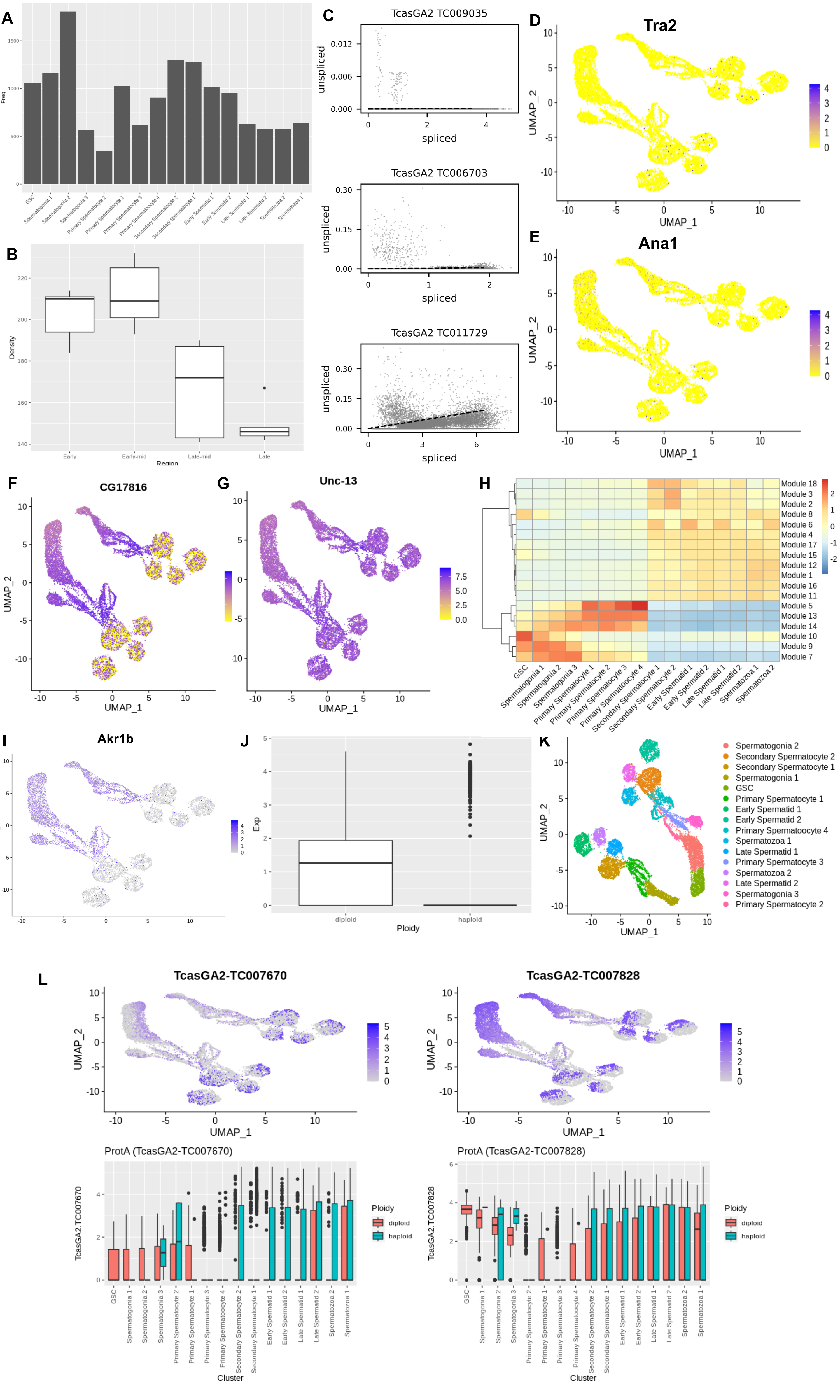

### SupplementaryFigure3.png

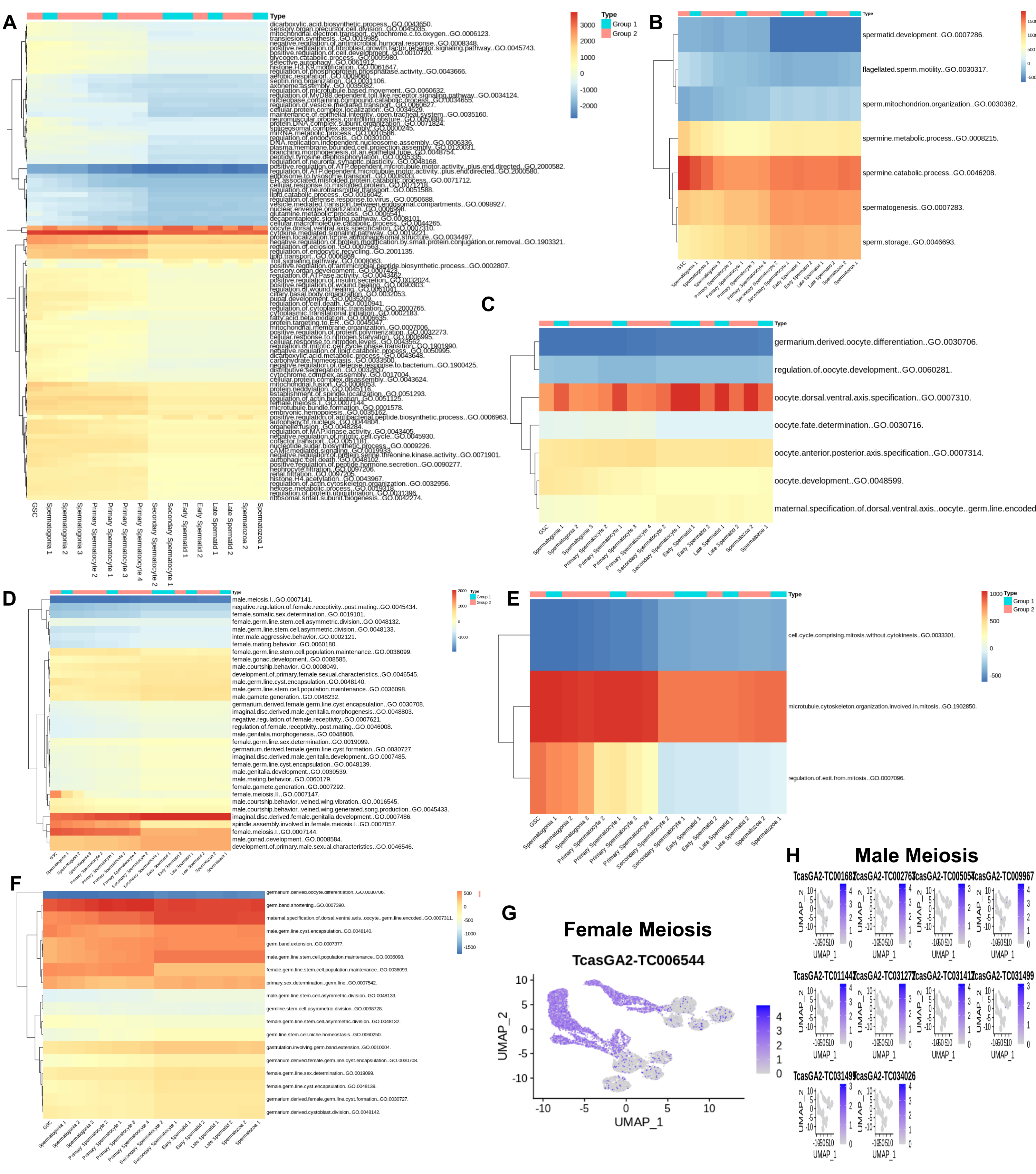

### SupplementaryFigure4.png

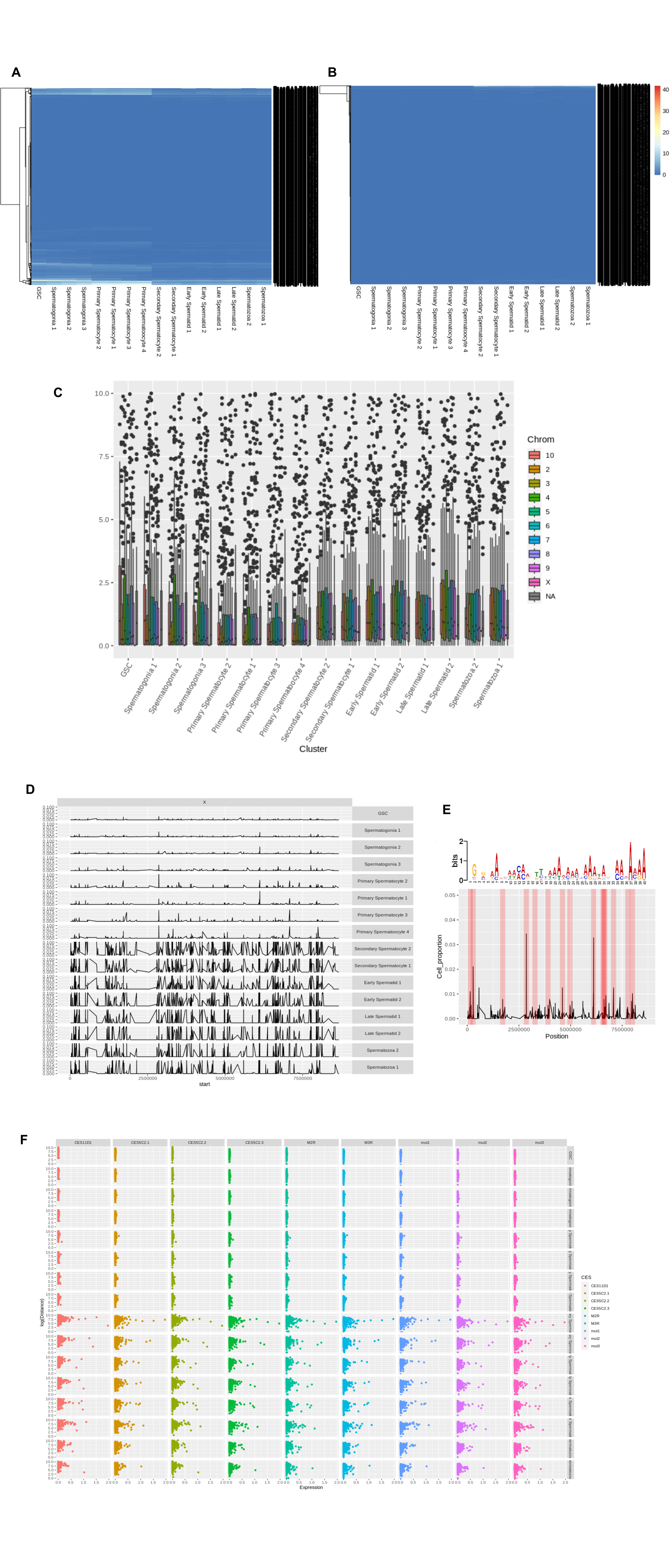
